# Predicted structure of the complete CPLANE complex reveals novel interactors and mechanisms of Jbts17 during ciliogenesis

**DOI:** 10.64898/2026.10.05.756796

**Authors:** Tynan Gardner, Anastasia Prolomova, Chanjae Lee, Edward M. Marcotte, John B. Wallingford

## Abstract

Cilia are highly conserved organelles that use hundreds of unique proteins to drive extracellular motility and serve as signaling hubs for eukaryotic cells. The Ciliogenesis and Planar Polarity Effector (CPLANE) protein complex controls basal body docking during early ciliogenesis and recruits the intraflagellar transport (IFT) proteins to the base of growing cilia. Leveraging recent advantages in deep learning neural networks and high-performance computing, we predict the structure of JBTS17, the largest CPLANE subunit, and the tertiary structure of the complete CPLANE complex, revealing direct contacts between JBTS17 and three other CPLANE members. We then used this model for an *in silico* protein-protein interaction screen that identified several novel JBTS17 interactors. Experiments in *Xenopus* confirmed a functional link *in vivo* between Jbts17 and the Spinocerebellar ataxia-associated protein TTBK2. The data provide new insights into mechanisms of CPLANE function during ciliogenesis and demonstrate the power of modeling protein complexes for the discovery of physical and functional protein interactions in cell homeostasis and disease.

## Introduction

Cilia are microtubule-based organelles that are vital for vertebrate development. Mutations that disrupt cilia motility, signaling, or biogenesis result in a pleiotropic set of human diseases called ciliopathies, which have a diverse set of symptoms ranging from kidney cysts, obesity, and craniofacial disorders to respiratory insufficiency and *situs inversus* (Wallmeier *et al*., 2020; Hilgendorf *et al*., 2024). Around 10% of the mammalian genome encodes ciliary proteins (Van Sciver and Caspary, 2024), and while null mutations to many of these genes cause embryonic lethality, hypomorphic alleles often cause lifelong disease in patients (Fliegauf *et al*., 2007). Understanding the disease etiology of these alleles requires an acute understanding of their respective proteins’ structure, interactions, and molecular mechanisms, which are often difficult problems to solve at scale using traditional cell biology techniques.

The five-member CPLANE (Ciliogenesis and Planar Polarity Effector) complex is one such set of proteins that are indispensable for ciliogenesis in vertebrates and causes ciliopathies when perturbed (Adler and Wallingford, 2017; Martín-Salazar and Valverde, 2022). CPLANE, which localizes to the transition zone of mature cilia (Figures 1A-A’), drives early migration of ciliary basal bodies from the cytoplasm to the apical surface, coordinates apical actin, and recruits a subset of the intraflagellar transport (IFT) proteins (Park *et al*., 2006, 2008; Brooks and Wallingford, 2012; Toriyama *et al*., 2016). The complex is composed of RSG1, FUZ, INTU, WDPCP, and JBTS17, which assemble at basal bodies to coordinate these processes (Toriyama *et al*., 2016).

**Figure 1.**
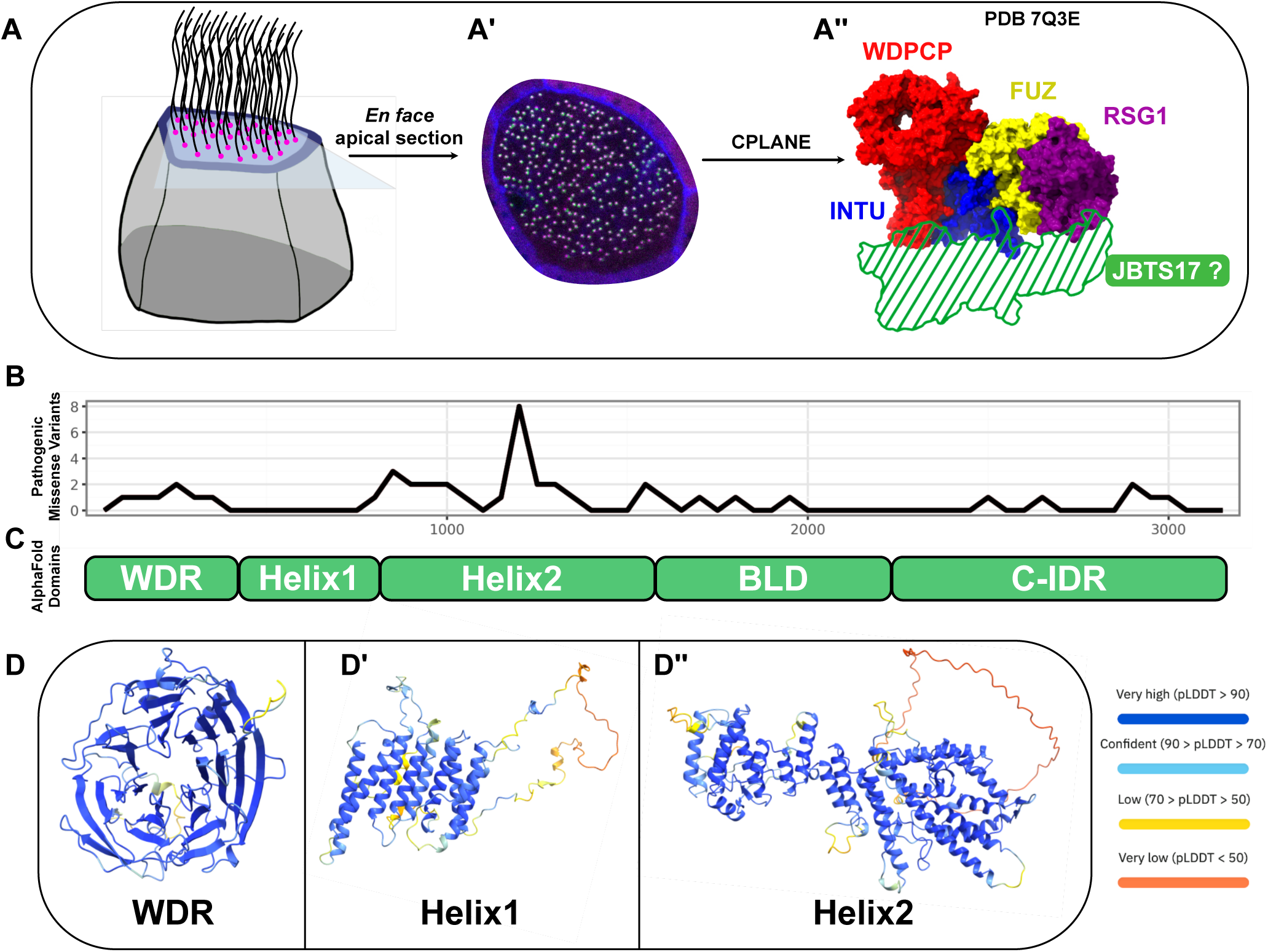
AlphaFold predicts the structure of JBTS17. (A) Schematic representation of a multiciliated cell and CPLANE localization. Translucent plane indicates the apical surface. (A’) Representative image of CPLANE localization in multiciliated cells. GFP-RSG1 accumulates at basal bodies. (A’’) Schematic of the incomplete CPLANE complex structure (PDB 7Q3E). JBTS17 lacks a solved or predicted structure. (B) Distribution of pathogenic missense JBTS17 variants discovered in human patients. (C) Schematic of the predicted structural organization of JBTS17 based on the AlphaFold2 model. Labeled regions correspond to the following residues: WD40 repeat-containing region (WDR), JBTS17^1-425^; first alpha-helical bundle (Helix1), JBTS17^426-818^; second alpha-helical bundle (Helix2), JBTS17^819-1569^; basal body-localizing domain (BLD), JBTS17^1570-2318^; and C-terminal intrinsically disordered region (C-IDR), JBTS17^2319-3197^. (D-D’’) AlphaFold2-predicted structures of selected JBTS17 fragments. Residues are color-coded by predicted Local Distance Difference Test (pLDDT) scores.

The disease etiology of JBTS17, CPLANE’s largest subunit at over 3,100 amino acids, remains particularly enigmatic. Variants in *JBTS17* can cause Joubert Syndrome, Oral-Facial-Digital Syndrome, and Meckel Gruber Syndrome, and over 60 variants have been clinically associated with these ciliopathies (Srour *et al*., 2012; Shaheen *et al*., 2013; Lopez *et al*., 2014; Romani *et al*., 2015). Despite its clinical significance, JBTS17 has proven harder to study than its four partners. For example, CryoEM structures have recently been solved for the other four subunits (Langousis *et al*., 2022), yet the tertiary structure of JBTS17 as well as its quaternary structure within CPLANE remain unknown (Figure 1A’’). Likewise, the cellular functions of Jbts17 are incompletely understood. In *Xenopus*, it is required to recruit the IFT-A2 proteins and the CPLANE subunits Rsg1, Wdpcp, and Intu to basal bodies (Toriyama *et al*., 2016). In mice, it is required for the localization of transition zone proteins NPHP1 and CEP290 to the transition zone in primary cilia (Damerla *et al*., 2015). However, the molecular mechanism of how it recruits these proteins to cilia remains unknown.

Advances in deep learning neural networks have enabled protein structure prediction at scale using AlphaFold (Jumper *et al*., 2021). This computational approach offers a new, scalable way to predict and explore protein-protein interactions from sequence and to understand potential structure-function relationships of patient alleles. While these algorithms are powerful, with databases beginning to accumulate predicted protein-protein interactions (Humphreys *et al*., 2021; Schmid and Walter, 2025), there are a growing number of efforts to follow up focused AlphaFold screens with *in vivo* validation, demonstrating the utility of these tools in the context of cell and developmental biology (McCafferty *et al*., 2022; van Breugel *et al*., 2022; Lim *et al*., 2023; Boegholm *et al*., 2024; Lee *et al*., 2024; Kawaguchi *et al*., 2025; Guo *et al*., 2026; Zhang *et al*., 2026). Furthermore, the AlphaFold database does not include full structures for monomeric proteins larger than 2,700 amino acids (Fleming *et al*., 2025). Because newer models and screens commonly follow a similar cutoff, JBTS17 and other large proteins have been omitted from most analyses (Schweke *et al*., 2024; Zhang *et al*., 2025; Han *et al*., 2026).

Here, we present the predicted structures for JBTS17 and the complete CPLANE complex. This complete model predicts a full set of interfaces between these subunits and helps explain the disease etiology of a clinically identified RSG1 allele. By taking advantage of high-performance computing, we applied the deep-learning protein structure prediction model AlphaFold2 at scale to systematically predict protein-protein interactions between JBTS17 and other proteins implicated in ciliogenesis. In addition to recapitulating known interactions, this screen predicted new links to basal body and distal appendage proteins, to axonemal proteins and to the apical actin network of multiciliated cells. We validated these findings with *in vivo* protein localization and recruitment assays in vertebrate embryos and identified a functional link to the Spinocerebellar Ataxia-associated protein TTBK2.

## Results and Discussion

### AlphaFold2 Predicts the Structure of JBTS17

Known pathogenic missense variants have been found throughout the length of JBTS17, with a prominent cluster around residues 800–1500 (Figure 1B). Despite the apparent functional importance of regions throughout the protein, InterPro lists only two predicted WD40 repeat-like domains at JBTS17^22–150^ and JBTS17^255–342^, with no annotations across the remainder of the protein (Blum *et al*., 2025). Moreover, because JBTS17 surpasses the 2,700 amino acid size threshold for the AlphaFold2 database (Fleming *et al*., 2025), no full-length predicted structure is currently available. By maximizing AlphaFold’s VRAM allocation on NVIDIA A100 GPUs, we were able to model protein monomers and multimer complexes of up to 8,000 amino acids and used this pipeline to model full-length human JBTS17.

Our model predicts that JBTS17 contains two core regions connected by a flexible linker, followed by a large C-terminal intrinsically disordered region (Figure 1C). The first high-confidence region contains a beta-propeller WDR domain connected to 9 bundled alpha helices (Figure 1D, 1D’). After an 18 amino acid flexible linker, the model contains a second bundle of 19 alpha helices (Figure 1D’’). The C-terminal 1,627 amino acids (JBTS17^1570–3197^) were predicted to be mostly disordered with a few low-confidence alpha helices.

Our model’s overall pTM score was 0.44 (Supplemental Figure 1A), which is below the usual confidence thresholds, but we note that the large region of intrinsic disorder is expected to reduce AlphaFold scores and interaction metrics (Banhos Danneskiold-Samsøe *et al*., 2024; Bret *et al*., 2024; Homma *et al*., 2024; Lee *et al*., 2024; Martin, 2024; Pándy-Szekeres *et al*., 2024; Dunbrack, 2025; Mitic *et al*., 2025). We therefore recomputed the model with only the high-confidence N-terminal region of the protein, omitting the C-terminal Intrinsically Disordered Region (IDR). Omitting this portion resulted in a higher pTM score of 0.61 and high confidence regional PAE values for individual residues in the WDR and alpha helical regions (Supplemental Figure 1B).

The predicted combination of a WDR beta-propeller adjacent to bundled alpha helices is reminiscent of the WDR-TPR domain combination observed in vesicle-trafficking proteins such as COP-I/II, Clathrin, SEC31, and several IFT proteins (Jékely and Arendt, 2006; van Dam *et al*., 2013; Field and Rout, 2022; McCafferty *et al*., 2022).

Comparison of JBTS17 with representative proteins containing this architecture further illustrates the similarity in their overall domain organization (Supplemental Figure 1E-J). This structure is interesting because it suggests a common protocoatomer evolutionary ancestor for these proteins, and hints at potential vesicle-related activity of JBTS17.

Accordingly, the CPLANE subunits INTU, FUZ, and WDPCP have also been shown to bind immobilized phosphoinositides (Langousis *et al*., 2022), and the first ∼800 amino acids of our JBTS17 model show a striking structural similarity to those of full-length WDPCP (Supplemental Figure 1K). Our model is therefore consistent with the hypothesis that CPLANE may bind vesicles.

### AlphaFold2 Models the Structure of the Complete CPLANE Complex

Using the same RAM allocation strategy to increase model token limits, we next modeled the full five-membered CPLANE complex with full length JBTS17 (pTM 0.64, ipTM 0.70; Supplemental Figure 1). Similar to the single-chain JBTS17 model, recomputing the model with the IDR-trimmed sequence JBTS17^1–1768^ resulted in higher confidence scores (pTM 0.75, ipTM 0.75; Figure 2A-B). Superposition of our predicted complex with the experimentally determined partial human CPLANE structure (PDB 7Q3D) showed close agreement in the organization of the shared WDPCP, INTU, and FUZ subunits (Supplemental Figure 2A).

**Figure 2.**
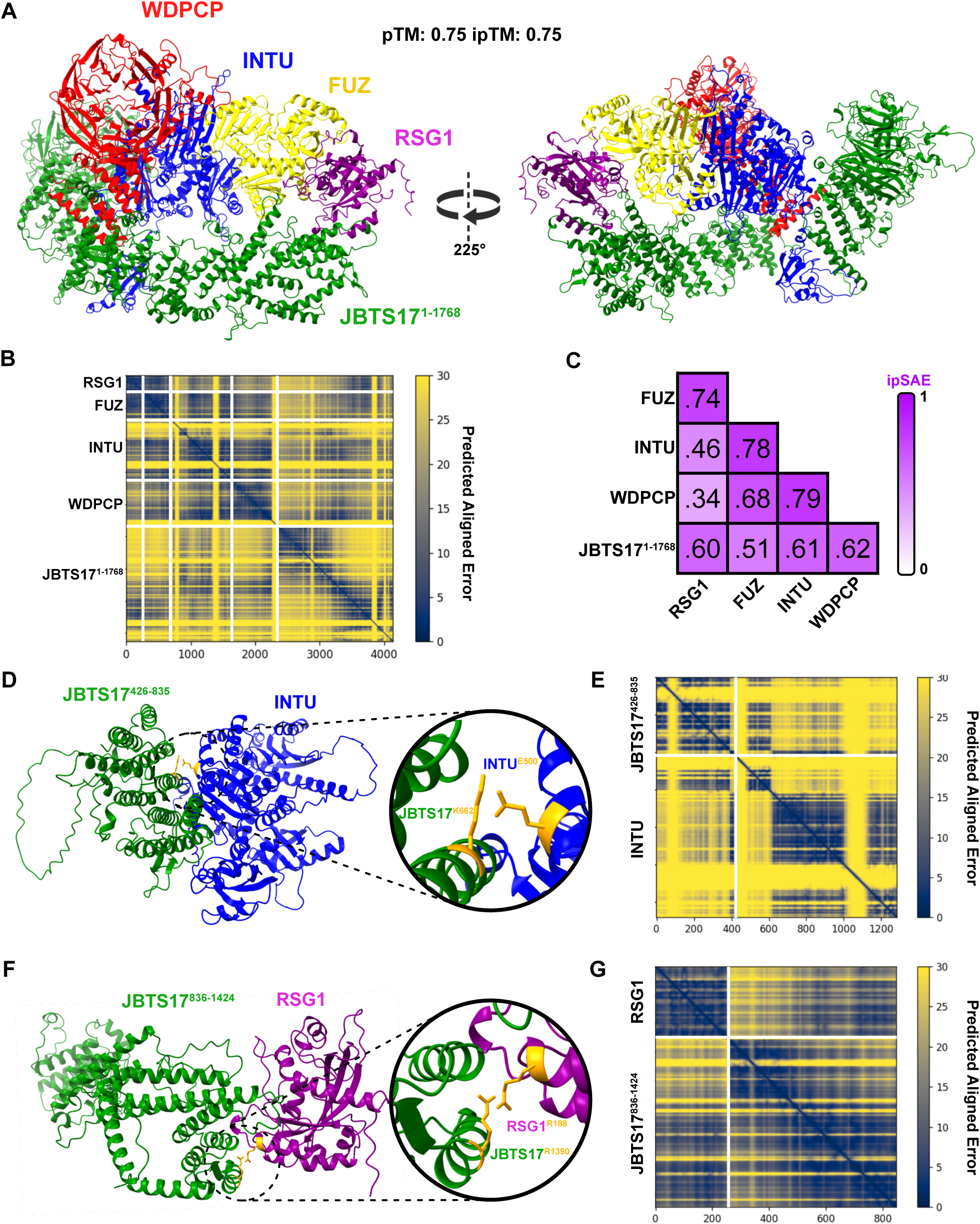
AlphaFold models the full CPLANE structure and a disease-implicated interface. (A) AlphaFold2-predicted structure of the full five-membered CPLANE complex with JBTS17^1-1768^, with the C-terminal intrinsically disordered region omitted. For clarity of presentation, small regions of low order are hidden. (B) Predicted Aligned Error (PAE) plot for the full CPLANE complex. (C) Pairwise ipSAE score matrix for CPLANE subunits. (D) AlphaFold2-predicted heterodimer between JBTS17^426-835^ and INTU. INTU^E500^, a residue mutated in a patient variant (Toriyama *et al*., 2016), is highlighted in orange along with adjacent residue JBTS17^K662^ (inset). pTM = 0.52, ipTM = 0.65, ipSAE^max^ = 0.46 (E) PAE plot for the JBTS17^426-835^-INTU model. (F) AlphaFold2-predicted heterodimer between JBTS17^836-1424^ and RSG1. RSG1^R188^, a residue mutated in a patient variant (Vazquez *et al.,* 2025), is highlighted in orange along with adjacent residue JBTS17^R1390^ (inset). pTM = 0.78, ipTM = 0.78, ipSAE = 0.54 (G) PAE plot for the JBTS17^836-1424^-RSG1 model.

We further analyzed this model using the alternative metric ipSAE^max^, which reduces the influence of disordered and non-interacting regions and has shown improved discrimination of true and false protein interactions compared with ipTM (Dunbrack, 2025; Mitic *et al*., 2025), to generate pairwise scores for individual CPLANE subunit pairs (Figure 2C). Together with the full multimer model, these data predict that JBTS17 binds RSG1, INTU, and WDPCP but not FUZ. Conversely, FUZ is predicted to bind RSG1, INTU, and WDPCP but not JBTS17. This finding is consistent with the fact that JBTS17 recruits all CPLANE subunits except FUZ to basal bodies (Toriyama *et al*., 2016). Finally, RSG1 is predicted to reside at the complex periphery, interacting with JBTS17 and FUZ but not WDPCP or INTU (Figure 2A).

### CPLANE Model Provides Insight into JBTS17 Interfaces Implicated in Ciliopathies

Variants in the genes encoding all CPLANE subunits have been shown to cause human ciliopathy (Martín-Salazar and Valverde, 2022), and our predicted model provides testable predictions that should inform disease etiology. For example, ClinVar reports numerous JBTS17 missense variants of uncertain significance (VUS), 19 of which map within 4 Å of predicted interfaces between JBTS17 and other CPLANE subunits in our model (Supplemental Figure 2B-E). These interface-associated VUS occur at predicted contacts with INTU, WDPCP, FUZ, and RSG1 and therefore provide testable candidates for investigating how these JBTS17 variants may affect CPLANE complex assembly or function. Conversely, our model predicts that INTU E500 lies at the interface with JBTS17 (Figure 2D-E), suggesting a possible molecular basis for the impaired recruitment of IFT-A subunits previously observed for the ciliopathy-associated INTU^E500A^ variant in *Xenopus* (Toriyama *et al*., 2016). Notably, INTU^E500A^ was stably expressed *in vivo* in that study, suggesting that this phenotype does not result from gross destabilization of INTU.

Of particular interest to us was the interface between JBTS17 and RSG1, the latter of which was recently linked directly to the ciliopathy Oral-Facial-Digital Syndrome in patients (Vazquez *et al*., 2024). In that study, one patient presented with cleft palate, choanal stenosis, and kidney cysts and was found to harbor the missense variant *RSG1^R188W^*. This altered residue mapped to the external surface of the experimentally determined partial CPLANE structure (Langousis *et al*., 2022), leaving its molecular impact unclear.

Our model of the complete CPLANE complex suggested that RSG1 R188 lies directly at the interface between RSG1 and JBTS17. To investigate this interface more closely, we specifically modeled the interaction between RSG1 and JBTS17^836–1424^, a smaller fragment of JBTS17 that contained that interacting region and produced higher confidence scores (pTM 0.78, ipTM 0.78, ipSAE^max^ 0.54). In this model, RSG1 R188 is positioned adjacent to JBTS17 R1390 (Figure 2F-G). Although arginine residues can participate in contacts at protein-protein interfaces (Neves *et al*., 2012), experimental studies will be required to determine whether these residues interact directly and how the R188W substitution affects this interface. Importantly, RSG1^R188W^ produces stable, soluble protein that localizes to basal bodies *in vivo* (Vazquez *et al*., 2024), arguing against gross protein destabilization as an explanation for its pathogenic effect.

Consistent with these in vivo observations, computational analyses predicted limited effects on protein stability for both variants. Missense3D (Ittisoponpisan et al., 2019) classified RSG1R188W as structurally benign, while DynaMut2 (Rodrigues et al., 2021) predicted only a small destabilizing effect (ΔG = −0.22 kcal/mol) and FoldX (Delgado et al., 2019) predicted a modest destabilizing effect (ΔΔG = +0.90 kcal/mol). INTU was likewise predicted to have little effect on protein stability by DynaMut2 (ΔΔG = +0.04 kcal/mol, stabilizing) and FoldX (ΔΔG = +0.19 kcal/mol), although interpretation of structure-based predictions at this residue is limited by its relatively low AlphaFold confidence (pLDDT = 54.25), which also prevented evaluation by Missense3D. Together with the in vivo stability of both mutant proteins, these results support investigating whether these ciliopathy-associated substitutions instead alter CPLANE protein-protein interactions without substantially disrupting the stability of the individual subunits.

### An AlphaFold2 Multimer Screen Predicts Novel JBTS17 Protein Interactors

We next sought to extend our analysis of JBTS17’s interactions. By dividing human JBTS17 into 5 fragments each less than 1,000 amino acids long, we performed a high-throughput protein-protein interaction screen *in silico* against a selected set of 448 target human proteins known to be in cilia or involved in ciliogenesis. The five JBTS17 bait fragments comprised the WDR beta-propeller, the first bundle of alpha helices and linker residues, the second bundle of alpha helices, the basal body-localizing domain (BLD), which was previously shown to be sufficient for basal body localization of JBTS17 (Toriyama *et al*., 2016), and the remainder of the C-terminal IDR (Figure 3C). The target set was created by combining the intersections of 3 datasets: the CiliaCarta (van Dam *et al*., 2019), targets of the ciliogenic transcription factor Rfx2 (Chung *et al*., 2014), and hits from previous CPLANE Affinity Purification Mass Spectrometry experiments (Toriyama *et al*., 2016), plus thirty-eight manual additions known to be ciliogenic but not found in the above datasets (Xu *et al*., 2026) (Figure 3A).

**Figure 3.**
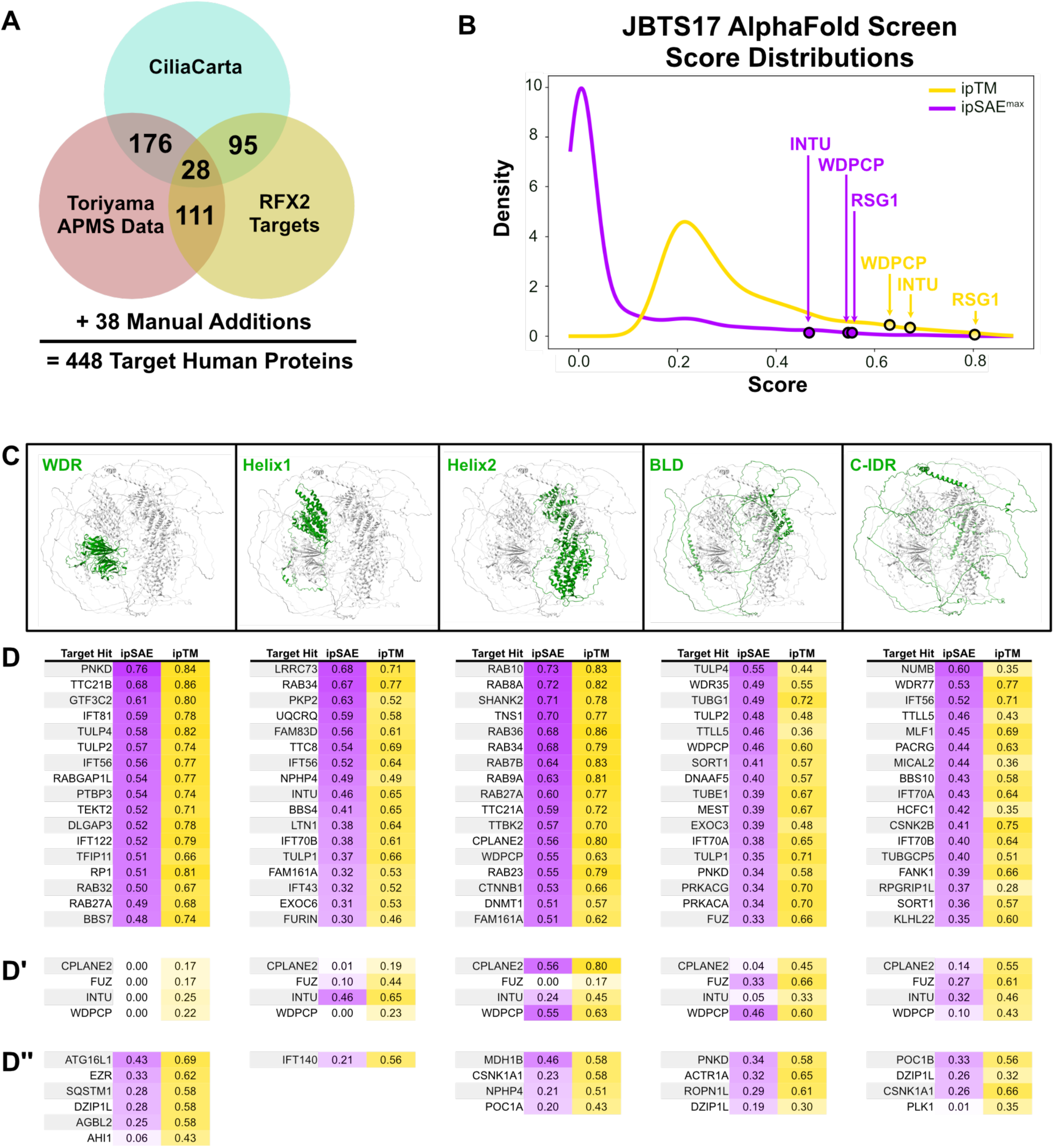
An AlphaFold screen predicts interactors of JBTS17 in silico. (A) Venn Diagram depicting AlphaFold screen target selection. Human proteins that were present in two or more shown datasets were selected, plus 38 manual additions for a total of 448 target human proteins. (B) Distribution of AlphaFold2 ipTM and ipSAE scores across all predicted models. Top scores for CPLANE members INTU, WDPCP, and RSG1 are highlighted with arrows. (C) AlphaFold2-predicted structure of full-length JBTS17 with bait fragments highlighted in green. (D) Top 17 hits for each bait fragment, sorted by ipSAE score. (D’) Scores of other CPLANE subunits for each bait fragment. (D’’) Other selected target hits that either scored highly via ipTM or were biologically interesting.

When combined with the five JBTS17 fragments, we obtained a full set of 2,240 possible combinations that were run using the structure prediction model ColabFold multimer (Mirdita *et al*., 2022). Predicted interactions were then scored both by AlphaFold’s default confidence score, ipTM, and also by ipSAE^max^. Target hits were sorted by ipSAE^max^, and a selection of top hits chosen for further analysis (Figure 3D). Scores for all tested combinations are presented in Supplemental Table 1.

As expected, most targets were not predicted to bind JBTS17, leading to a positively skewed distribution of both ipTM and ipSAE^max^ scores (Figure 3B). As an important positive control, CPLANE complex members RSG1, INTU, and WDPCP all ranked highly with both scores (Figure 3B and 3D’), recapitulating the known CPLANE subunit interactions determined by proteomics (Toriyama *et al*., 2016; Vazquez *et al*., 2024).

These known positive interactions between CPLANE subunits show higher pairwise ipTM versus ipSAE^max^ scores (Figure 3D’). However, ipSAE^max^ scores across the entire screen extend closer to zero than ipTM scores, which are largely compressed above ∼0.2, resulting in a broader observed range of ipSAEmax scores and greater numerical separation between the highest-scoring interactions and the bulk of screened proteins (Figure 3B).

In addition to recruiting three other CPLANE subunits, JBTS17 is required for the localization of the IFT-A2 subcomplex to basal bodies (Toriyama *et al*., 2016). The three IFT-A2 proteins score highly with various JBTS17 fragments in our screen. TTC21B (IFT139) is the second highest hit for the JBTS17 WDR fragment. WDR35 (IFT121) is the second highest hit for the JBTS17 BLD fragment, albeit with lower scores of ipSAE^max^ 0.49 and ipTM 0.55. IFT43 scores lower still, with ipSAE^max^ 0.32 and ipTM 0.52 for JBTS17 fragment Helix1 (Figure 3D). JBTS17 is not required for the recruitment of IFT-B or IFT-A1 proteins (Toriyama *et al*., 2016), yet several of these proteins scored highly in our screen, including IFT56, IFT70A/B, IFT81, and IFT122. As recruitment dependency does not necessarily imply or exclude physical interaction, these predictions raise the possibility that some IFT proteins contact JBTS17 without depending on it for recruitment. Alternatively, these predicted interactions could reflect contacts that occur in the context of assembled IFT/CPLANE complexes. Experimental studies will be required to determine whether these predicted interactions occur in vivo.

### Candidate JBTS17 Interactors Suggest Diverse Roles for Jbts17 in MCCs

We selected several top candidates based on ipSAE^max^ and ipTM scores for further analysis *in vivo* using *Xenopus laevis* multiciliated cells (MCCs), a powerful platform for investigating ciliary cell biology (Walentek and Quigley, 2017). To determine their subcellular localization, we injected *Xenopus* embryos with mRNA encoding GFP-tagged candidate proteins and imaged their subcellular localization in the mucociliary epithelium by confocal microscopy. Predicted structures of the candidate JBTS17 interactions discussed below that are not shown elsewhere are presented in Supplemental Figure 4. As expected based on the localization of Jbts17 and the CPLANE complex, many top candidates localized at or near basal bodies, including Ttbk2, Csnk1a1, Ahi1, Plk1, Poc1a, Poc1b, and Bbs4 (Figure 4A-B and Supplemental Figure 3A-E). While Csnk1a1 has been shown to localize to the basal body in mouse primary cilia (Chong *et al*., 2015), we observed Csnk1a1 localization to both basal bodies and the apical plasma membrane in the *Xenopus* mucociliary epithelium (Figure 4B).

**Figure 4.**
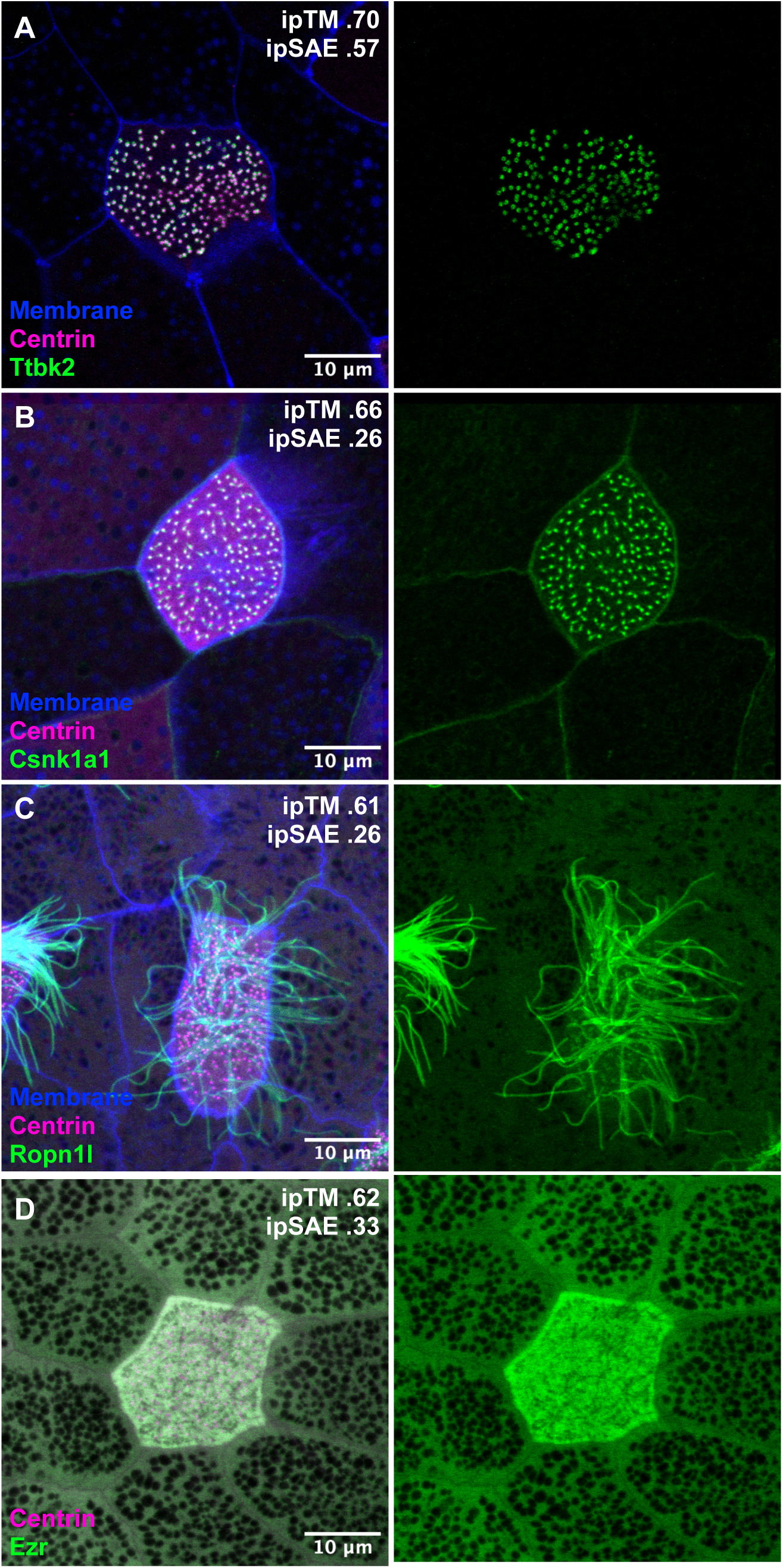
Predicted JBTS17 interactors localize throughout multiciliated cells. (A) GFP-tagged Ttbk2 localizes near basal bodies in an MCC. (B) GFP-tagged Csnk1a1 localizes near basal bodies in an MCC and membranes throughout the epithelium. (C) GFP-tagged Ropn1l localizes to ciliary axonemes in an MCC. (D) GFP-tagged Ezrin localizes to the apical cap of an MCC. Centrin is visualized by co-expressed Centrin2-RFP (magenta), and membranes are visualized by co-expressed membrane-BFP (blue). Scale bars = 10 µm.

In addition to its role in ciliogenesis, CPLANE is implicated in the assembly of the specialized apical actin network that characterizes MCCs (Park *et al*., 2006; Yasunaga *et al*., 2015; Sharma *et al*., 2024). We were interested, then, to find that Ezrin (EZR) scored highly by ipTM with JBTS17’s WDR fragment. As found previously in mammalian MCCs (Huang *et al*., 2003; Yasunaga *et al*., 2022), we found EZR strongly localized to the MCC apical actin network (Figure 4D), suggesting that the Jbts17/Ezr interaction may be functionally significant.

In mature cilia, CPLANE resides at the transition zone (Vazquez *et al*., 2024), which controls the entry of axonemal proteins (Gonçalves and Pelletier, 2017). We were interested then to find that three candidate interactors, all previously identified as radial spoke proteins, localized to axonemes in MCCs: Ropn1l, Nme5, and Mdh1b (Figure 4C and Supplemental Figure 3F-G) (Gui *et al*., 2021; Walton *et al*., 2023; Leung *et al*., 2025). Because axonemal proteins are known to make physical contacts with transition zone proteins that gate them (Takao *et al*., 2017), these findings raise the possibility that CPLANE interacts with radial spoke proteins during their entry into motile cilia.

Actr1a, which encodes the dynactin subunit α-centractin/Arp1, was also ranked highly by ipTM and was of interest given established roles for dynactin in centrosome-associated trafficking and ciliogenesis (Dammermann and Merdes, 2002; Kodani *et al*., 2013). However, Actr1a showed diffuse localization throughout the cytoplasm of multiciliated cells with variable intensity across individual cells, without obvious enrichment at basal bodies (Supplemental Figure 3H).

Sqstm1 and Atg16l1, both autophagy related proteins, localized to bright punctae throughout the mucociliary epithelium (Supplemental Figure 3I-J). These candidates were of particular interest given previous findings that CPLANE proteins Fuz, Intu, and Wdpcp bind the phospholipid PI(3)P, a key regulator of autophagy initiation (Burman and Ktistakis, 2010; Marat and Haucke, 2016; Langousis *et al*., 2022). Furthermore, autophagic degradation controls the removal of Ofd1 (oral-facial-digital syndrome 1), a distal appendage protein linked to some of the same ciliopathies as CPLANE subunits, from centriollar satellite pools (Tang *et al*., 2013; Bruel *et al*., 2017).

Many Rab proteins scored highly in our screen, especially with the Helix2 fragment of JBTS17 (Figure 3D). Rabs and related small GTPases are vital during early ciliogenesis; in particular, the Rab11-Rabin8-Rab8 cascade drives transport of small Distal Appendage Vesicles to migrating basal bodies, and coordinate fusion of these vesicles into the larger Ciliary Vesicle (Nachury *et al*., 2007; Knödler *et al*., 2010; Westlake *et al*., 2011; Blacque *et al*., 2017). The two Rabs with the highest ipSAE^max^ in our screen, Rab8 and Rab10, localized to Golgi without strong signal near basal bodies in MCCs (Supplemental Figure 3K-L), despite the fact that these proteins localize to primary cilia (Yoshimura *et al*., 2007; Babbey *et al*., 2010). We noted that Rab10 also localized to secretory vesicles in the goblet cells of the mucociliary epithelium, which is interesting because the CPLANE subunit FUZ is required for exocytosis from these vesicles (Gray *et al*., 2009). We note, however, that due to the close structural similarity of the Rab family and potential redundancy between Rabs (Sato *et al*., 2014), we cannot rule out the possibility that these high scores are nonspecific and indicate binding to other related Rabs.

An important limitation of this approach is that AlphaFold-based interaction predictions do not provide information about the spatial or temporal dynamics of an interaction in vivo. Thus, our models cannot distinguish whether candidate proteins remain associated with JBTS17 after recruitment to basal bodies or cilia, interact only transiently during trafficking or assembly, or interact in a different cellular context. Future biochemical and in vivo imaging studies will be required to establish whether these predicted interactions occur and to determine their dynamics within cells.

### Jbts17 is Required for Ttbk2 Localization to Basal Bodies

To test if the modeled physical interactions reflect functional interactions *in vivo*, we explored the link between JBTS17 and TTBK2, a well-known distal appendage protein that recruits RSG1 to basal bodies and has a similar localization to JBTS17 (Goetz *et al*., 2012; Agbu *et al*., 2018). TTBK2 comprises a core kinase domain, a long flexible disordered region, and a small alpha helix at its C-terminal that is predicted to bind JBTS17 at the latter’s amino acids 819–1569 (Figure 5A-B). In the disease allele TTBK2^SCA11^, the C-terminal half of the protein including this helix is missing; this truncated protein is unable to localize to the basal body, and patients present with Spinocerebellar ataxia type 11 (Goetz *et al*., 2012; Bowie *et al*., 2018).

**Figure 5.**
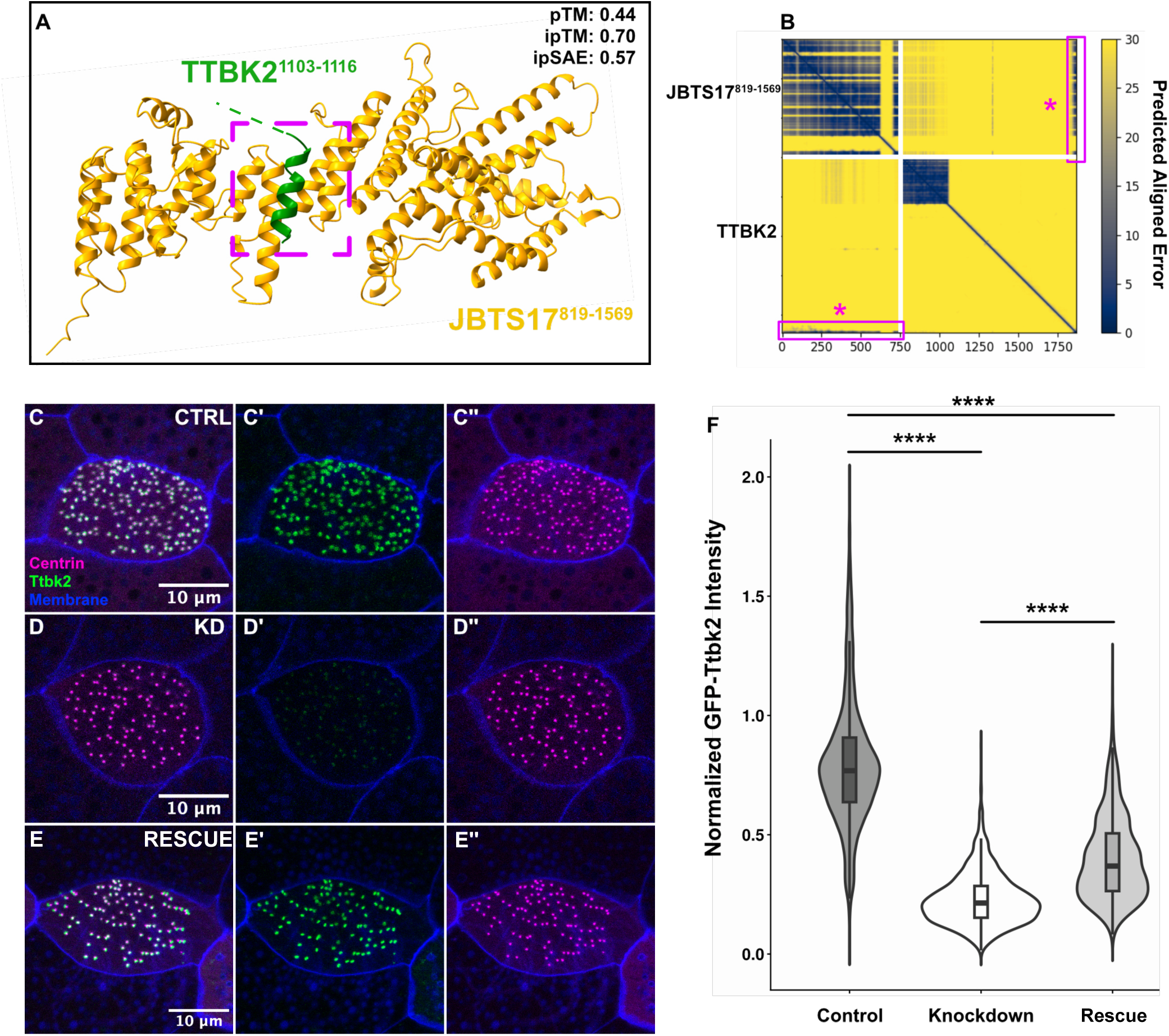
Known ciliogenesis regulator and transition zone protein Ttbk2 is localized to basal bodies by Jbts17. (A) AlphaFold2-predicted structure of JBTS17^819-1569^-TTBK2 heterodimer. For clarity of presentation, the N-terminal kinase region and intrinsically disordered region of TTBK2 are hidden. (B) PAE plot for the JBTS17^819-1569^-TTBK2 model. Data corresponding to the interacting regions are indicated with magenta boxes and asterisks. (C-E″) Localization of GFP-tagged Ttbk2 to basal bodies, marked by Centrin-RFP, in control MCCs (C-C″), Jbts17 knockdown MCCs (D-D″), and Jbts17 knockdown MCCs co-expressing Myc-Jbts17 (E-E″). Jbts17 knockdown strongly reduces Ttbk2 localization to basal bodies, while Myc-Jbts17 expression results in partial restoration of Ttbk2 basal body localization. Scale bars = 10 µm. (F) Quantification of GFP-Ttbk2 localization to basal bodies, normalized to Centrin-RFP intensity and averaged within individual MCCs. Control vs. Jbts17 knockdown, p = 5.4e-14; Jbst17 knockdown vs. rescue, p = 4.5e-05; control vs. rescue, p = 1.2e-10 (pairwise two-tailed Mann–Whitney tests).

To test whether Ttbk2 localization was disrupted upon knockdown of Jbts17, we used a previously validated splice-blocking morpholino against *jbts17* (Toriyama *et al*., 2016). We co-expressed GFP-tagged Ttbk2 to visualize its localization, centrin-RFP to label basal bodies, and membrane-BFP to label cell boundaries. Jbts17 knockdown strongly reduced Ttbk2 localization at basal bodies (p = 5.4e-14) (Figure 5C-F). To confirm the specificity of this phenotype, we co-injected the Jbts17 morpholino with *myc-jbts17* mRNA, which partially rescued Ttbk2 localization to basal bodies (p = 4.5e-05) (Figure 5C-F). Together, our heterodimer model and this *in vivo* functional interaction provide a molecular model for TTBK2 recruitment by JBTS17 during ciliogenesis.

Together, our structural models of full-length JBTS17 and the complete CPLANE complex provide a framework for understanding CPLANE organization and generating testable hypotheses about the molecular consequences of human variants in JBTS17, RSG1, and INTU. Extending this structural framework, our *in silico* protein-protein interaction screen identified candidate interactions spanning basal body, apical actin, and axonemal biology, several of which we localized *in vivo* in *Xenopus* multiciliated cells. Most notably, the screen identified a novel functional link between Jbts17 and the kinase Ttbk2. Previous work placed TTBK2 upstream of INTU and RSG1 during ciliogenesis initiation (Agbu *et al*., 2018), whereas our finding that Jbts17 depletion reduces Ttbk2 basal body localization suggests additional functional interdependence between CPLANE and the centriole maturation machinery. Our model further predicts a potential direct interaction between the structured core of JBTS17 and the C-terminal alpha-helical region of TTBK2. Determining whether this and other predicted interactions occur directly *in vivo*, as well as their spatial and temporal dynamics, will require further biochemical and imaging studies. Thus, our computational approach provides a lightweight, high-throughput framework for generating experimentally testable hypotheses about CPLANE function, while our *in vivo* results reveal a previously unrecognized functional relationship between Jbts17 and Ttbk2 during ciliogenesis.

## Materials and Methods

### AlphaFold modeling and structural analysis

Structures of JBTS17, the CPLANE complex, and the protein-protein interaction screen were modeled using AlphaFold2 and AlphaFold-Multimer as implemented in a local ColabFold (v1.5) installation on Linux-based lab servers and the TACC Lonestar6 cluster, both equipped with NVIDIA A100 GPUs. For large models exceeding the default GPU memory allocation, the XLA memory configuration was modified using the XLA_PYTHON_CLIENT_PREALLOCATE=false, TF_FORCE_UNIFIED_MEMORY=true, and XLA_PYTHON_CLIENT_MEM_FRACTION=3.2 environment variables to increase the fraction of memory available to AlphaFold2. Multimer sequence alignments were generated via MMseqs2 server calls. All models were generated with five recycles and without Amber relaxation. Human protein sequences were obtained from UniProt.

To mitigate the impact of intrinsic disorder on multimer prediction performance, N- and C-terminal intrinsically disordered regions (IDRs) were trimmed from all protein sequences based on UniProt annotations (The UniProt Consortium, 2025), while internal IDRs were retained to preserve flexibility and distance between separate ordered domains. Previous studies have shown that intrinsic disorder in proteins can attenuate the effectiveness of AlphaFold-Multimer models, disproportionately lowering interaction scores among positively interacting proteins and significantly increasing model runtime and memory usage (Banhos Danneskiold-Samsøe *et al*., 2024; Bret *et al*., 2024; Homma *et al*., 2024; Lee *et al*., 2024; Martin, 2024; Pándy-Szekeres *et al*., 2024; Dunbrack, 2025; Mitic *et al*., 2025).

For interaction screening, five JBTS17 fragments were used as baits. Target proteins were selected based on presence in at least two of the following datasets: Rfx2 transcriptional targets (Chung *et al*., 2014), prior CPLANE affinity purification-mass spectrometry (AP-MS) datasets (Toriyama *et al*., 2016), and the CiliaCarta compendium (van Dam *et al*., 2019). Additional candidates, including IFT, BBsome, and Rab-associated proteins, as well as recently identified ciliary proteins WDR88, CCDC30, MORN2, and MIIP (Xu *et al*., 2026) were manually included. Predicted complexes were rescored using the ipSAE^max^ metric (Dunbrack, 2025), and structural models were visualized and analyzed in UCSF ChimeraX (Meng *et al*., 2023). Structural superpositions were performed using the ChimeraX Matchmaker tool.

Germline missense variants classified as variants of uncertain significance (VUS) for CPLANE1/JBTS17 were obtained from ClinVar using the GRCh38 assembly (accessed September 9, 2026). VUS positions were mapped onto the AlphaFold2-predicted CPLANE complex, and JBTS17 residues containing VUS were classified as interface-associated when any atom was located within 4 Å of another CPLANE subunit.

Contacts involving low-confidence/disordered regions omitted from the structural model visualization were excluded from this analysis.

### Missense variant stability analysis

The effects of RSG1^R188W^ and INTU^E500A^ on protein stability were evaluated using Missense3D (Ittisoponpisan *et al*., 2019), DynaMut2 (Rodrigues *et al*., 2021), and FoldX v5.1 (Delgado *et al*., 2019). AlphaFold-predicted monomer structures were used as inputs. Missense3D and DynaMut2 analyses were performed using their respective web servers with default parameters. For FoldX, structures were first optimized using the RepairPDB command, followed by mutation modeling using BuildModel with five modeling runs. FoldX ΔΔG values are reported as the mean across these runs. INTU E500 was excluded from Missense3D analysis due to insufficient local structural confidence.

### *Xenopus* embryo manipulations

All *Xenopus* experiments were conducted following the animal ethics guidelines of the University of Texas at Austin, protocol number AUP-2024-00130. Female adult *X. laevis* were induced to ovulate by injection of human chorionic gonadotropin. The following day, females were squeezed, and eggs were fertilized *in vitro* with homogenized testis. Embryos were dejellied in 1/3x Marc’s modified Ringer’s (MMR) with 3% (wt/vol) cysteine (pH 7.9) at the two-cell stage. All embryos were reared in 1/3x MMR solution until stage 26. For microinjections, embryos were placed in 2% Ficoll in 1/3 x MMR and injected using a glass needle, forceps, and an Oxford universal micromanipulator.

### Plasmids and cloning

Gene sequences for *Xenopus laevis* were obtained from Xenbase (www.xenbase.org) (Chu *et al*., 2025). Open reading frames were amplified from a *Xenopus* cDNA library by polymerase chain reaction and then inserted into a pCS10R vector containing a GFP tag. All constructs were verified by sequencing. The list of cloned genes is as follows: Ttbk2, Rab8a, Csnk1a1, Ropn1l, Ezr, Ahi1, Plk1, Poc1a, Poc1b, Bbs4, Nme5, Mdh1b, Actr1a, Atg16l1, Sqstm1, Rab10.

### mRNA and morpholino microinjections

Vectors were linearized by restriction digestion and capped mRNAs were synthesized using the mMESSAGE mMACHINE SP6 transcription kit (ThermoFisher Scientific, #AM1340). For knockdown experiments, we used the previously validated splice-blocking antisense morpholino oligonucleotide (MO) against *jbts17*, 5’- TCTTCTTGATCCACTTACTTTTCCC-3’ (GeneTools), which was previously shown to disrupt *jbts17* splicing by RT-PCR (Toriyama *et al*., 2016). *Xenopus* embryos were injected with mRNAs and/or MOs into the two ventral blastomeres at the four-cell stage to target the mucociliary epithelium. mRNAs for GFP-tagged candidates, Centrin-RFP, and membrane-BFP were injected at 100 pg per blastomere, and the *jbts17* morpholino was injected at 30 ng per blastomere. For rescue experiments, mRNA encoding Myc- Jbts17 was co-injected at 500 pg per blastomere.

### Live imaging and image analysis

For live imaging, stage 26 *Xenopus* embryos were mounted between two coverslips and submerged in 0.3x MMR and imaged using a Zeiss LSM700 confocal microscope using a Plan-Apochromat 63 × 1.4 NA oil DIC M27 immersion lens.

The fluorescence intensity of GFP-Ttbk2 at basal bodies was measured as previously described (Toriyama *et al*., 2016). Briefly, basal body regions of interest (ROIs) were identified in Fiji by first isolating the Centrin2-RFP channel. An intensity threshold was then applied, and particles within a defined size (0.05-0.75 µm²) and circularity (0.1-1.0) range were detected using the “Analyze Particles” function to generate ROIs corresponding to basal bodies. These ROIs were overlaid onto the original multichannel image, and fluorescence intensities were measured across the Centrin2-RFP and GFP- Ttbk2 channels. GFP-Ttbk2 fluorescence at each basal body was then normalized by dividing GFP intensity by the corresponding Centrin2-RFP intensity measured at the same ROI. Three independent injection experiments were performed, with control, Jbts17 knockdown, and rescue conditions generated from the same clutch of embryos in each experiment. For each condition, three embryos were imaged per experiment, with three multiciliated cells analyzed per embryo and at least 30 basal bodies quantified per cell.

### Statistical analysis and data visualization

Plots were generated using custom R scripts, utilizing the ggplot2 package (Wickham, 2016). Basal body measurements were averaged per cell prior to statistical analysis to account for non-independence. Fluorescence measurements from individual basal bodies were first averaged within each multiciliated cell, and cell-level means were analyzed using pairwise Wilcoxon-Mann-Whitney tests and Holm-adjusted p-values.Statistical analyses were performed in R using the base stats package.

## Acknowledgments

The authors gratefully acknowledge Anna Battenhouse and Biomedical Research Computing Facility for extensive assistance with computational analyses, and the Texas Advanced Computing Center at The University of Texas at Austin for providing computational resources that have contributed to the research results reported within this paper. Molecular graphics and analyses performed with UCSF ChimeraX, developed by the Resource for Biocomputing, Visualization, and Informatics at the University of California, San Francisco, with support from National Institutes of Health R01-GM129325 and the Office of Cyber Infrastructure and Computational Biology, National Institute of Allergy and Infectious Diseases. Research was funded by grants from the National Institute Of Dental & Craniofacial Research (F31 DE035794 to T.G.), National Institute of Child Health and Human Development (R01 HD085901 to J.B.W. and E.M.M.), National Institute of General Medical Sciences (R35GM122480 to E.M.M.), and the Welch Foundation (F-1515 to E.M.M.).

---

**Supplemental Figure 1.**
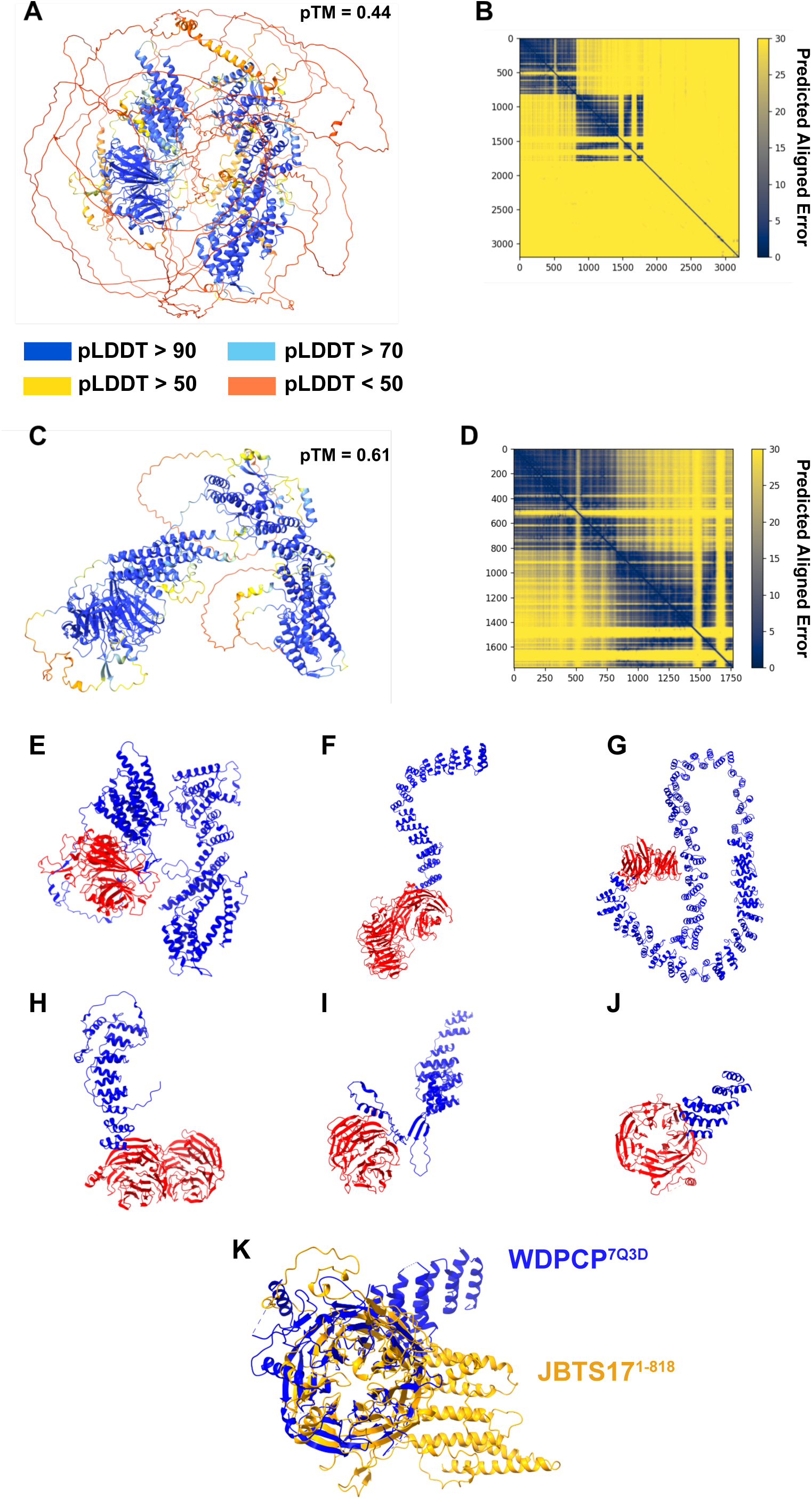
AlphaFold predicts the structure of JBTS17 and reveals similarities to other WDR–ɑ–helical proteins. (A) AlphaFold2-predicted structure of full length JBTS17. Residues are color-coded by predicted Local Distance Difference Test (pLDDT) scores. (B) PAE plot for the full length JBTS17 model. (C) AlphaFold2-predicted structure of IDR-trimmed JBTS17 (JBTS17^1–1768^). Residues are color-coded by predicted Local Distance Difference Test (pLDDT) scores. (D) PAE plot for the IDR-trimmed JBTS17 model. (E–J) Structural comparison of proteins containing WDR beta-propellers followed by alpha-helical regions: (E) human JBTS17, (F) human IFT140, (G) human clathrin heavy chain, (H) human COP1β, (I) yeast Sec31, and (J) human WDPCP. The experimentally determined structure is shown for IFT140 (PDB 8BBF) and WDPCP from the solved partial CPLANE structure (PDB 7Q3D), while AlphaFold-predicted structures are shown for the remaining proteins. (K) Structural superposition of the N-terminal WDR and alpha-helical regions of AlphaFold-predicted JBTS17 and experimentally determined human WDPCP (PDB 7Q3D), illustrating their similar domain architecture. Structures were superimposed using the ChimeraX Matchmaker tool.

**Supplemental Figure 2.**
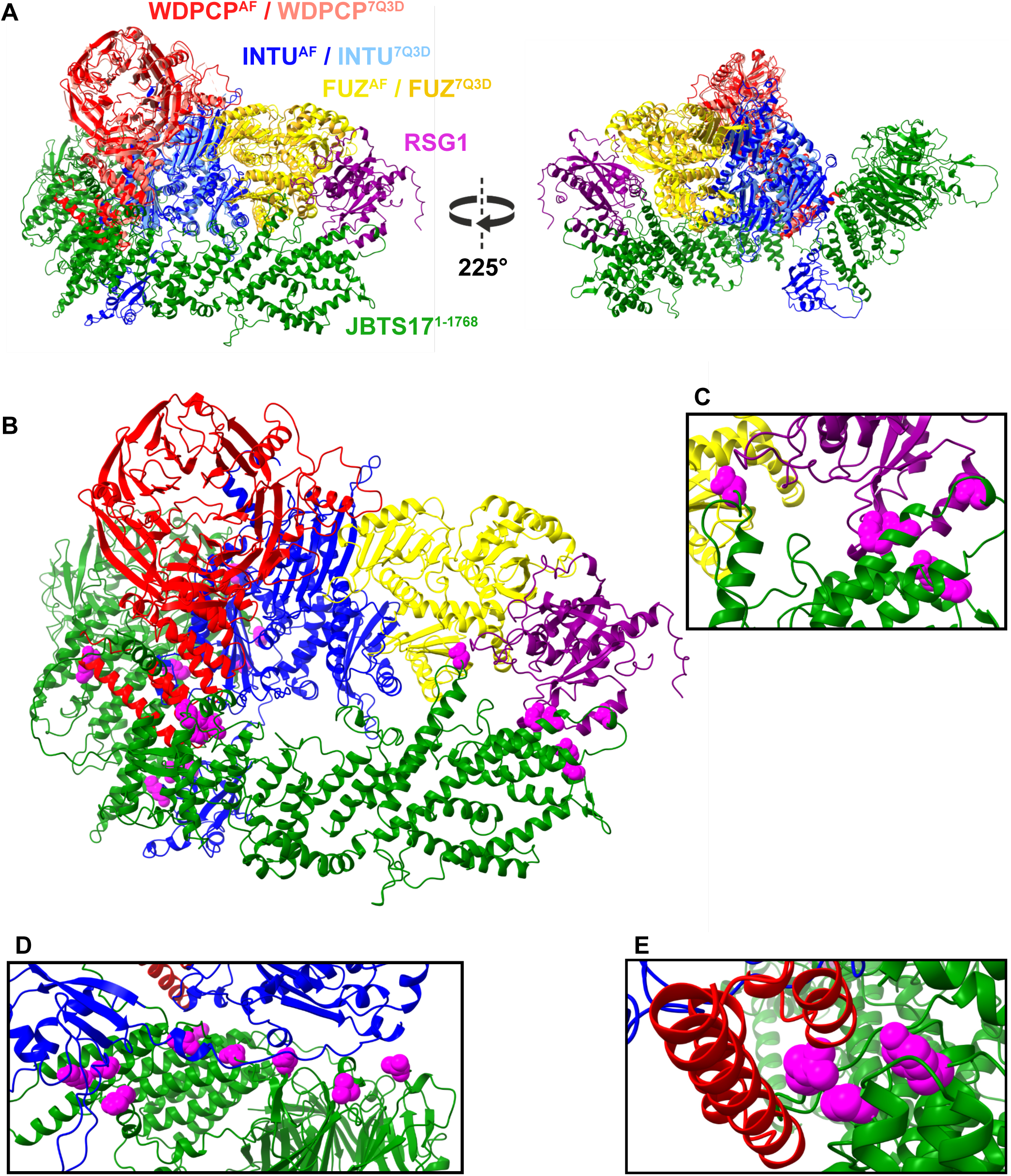
Structural comparison of the predicted CPLANE complex and mapping of JBTS17 variants of uncertain significance. (A) Superposition of the AlphaFold2-predicted five-membered CPLANE complex with the experimentally determined partial human CPLANE structure (PDB 7Q3D). The experimental structure contains human WDPCP, INTU, and FUZ, but not JBTS17 or RSG1. Corresponding subunits are shown in matching colors, with the experimental structure shown in lighter colors. The structures were superimposed using the ChimeraX Matchmaker tool. (B) Mapping of ClinVar missense variants of uncertain significance (VUS) onto the AlphaFold2-predicted CPLANE complex. JBTS17 residues containing VUS and located within 4 Å of another CPLANE subunit are highlighted in magenta and shown as spheres. Nineteen interface-associated VUS were identified after excluding contacts involving low-confidence/disordered regions omitted from the structural visualization. (C-E) Enlarged views of selected predicted JBTS17 interfaces containing ClinVar VUS. (C) VUS located at predicted interfaces with FUZ and RSG1. (D) VUS located at the predicted interface with INTU. (E) VUS located at the predicted interface with WDPCP. VUS-containing JBTS17 residues are shown in magenta.

**Supplemental Figure 3.**
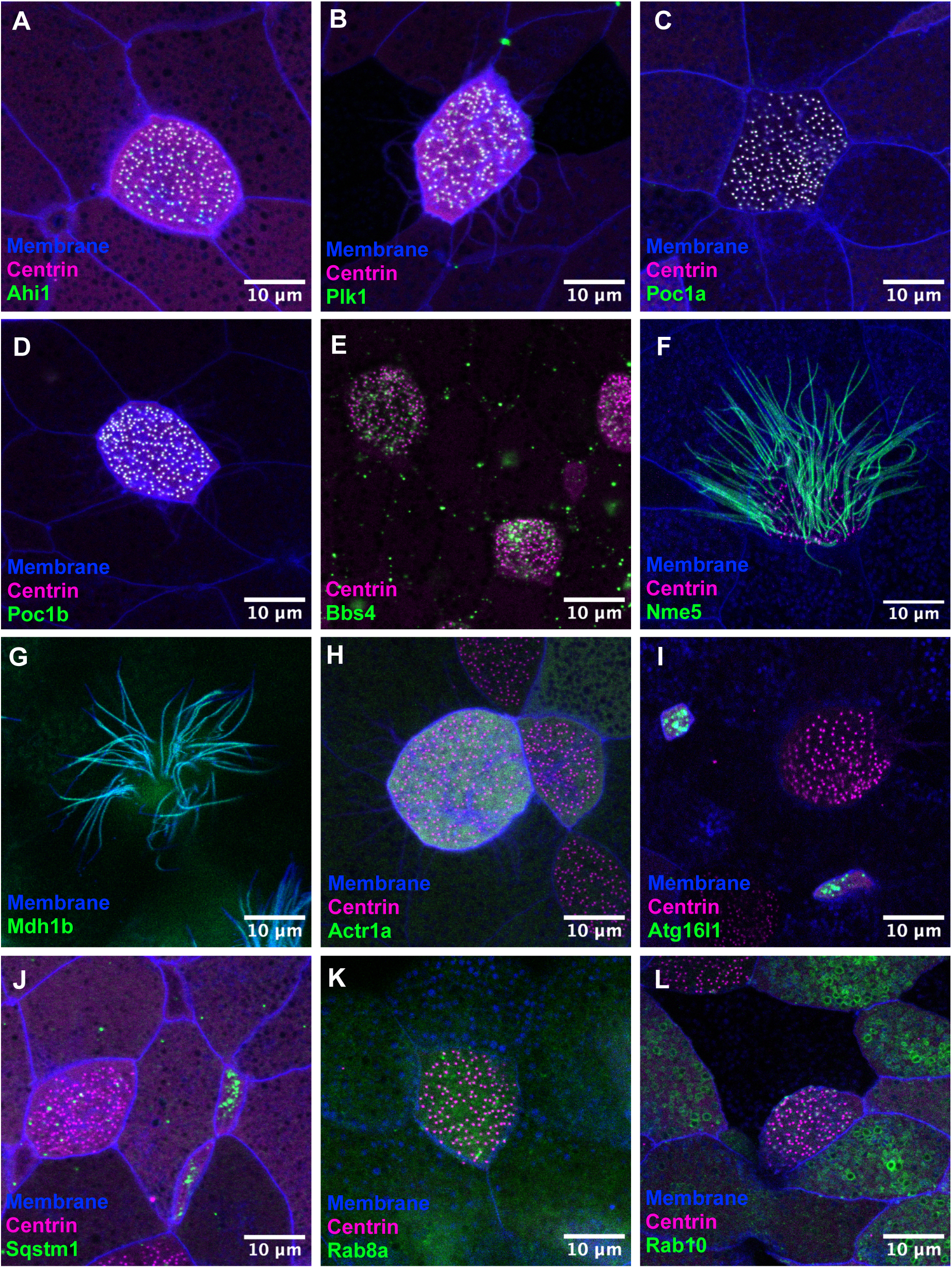
Localization of additional predicted JBTS17 interactors in multiciliated cells. (A-L) Representative images of *Xenopus* multiciliated cells expressing GFP-tagged candidate interactors. Cell membranes and basal bodies are marked by membrane-BFP and Centrin2-RFP, respectively. Scale bars = 10 µm.

**Supplemental Figure 4.**
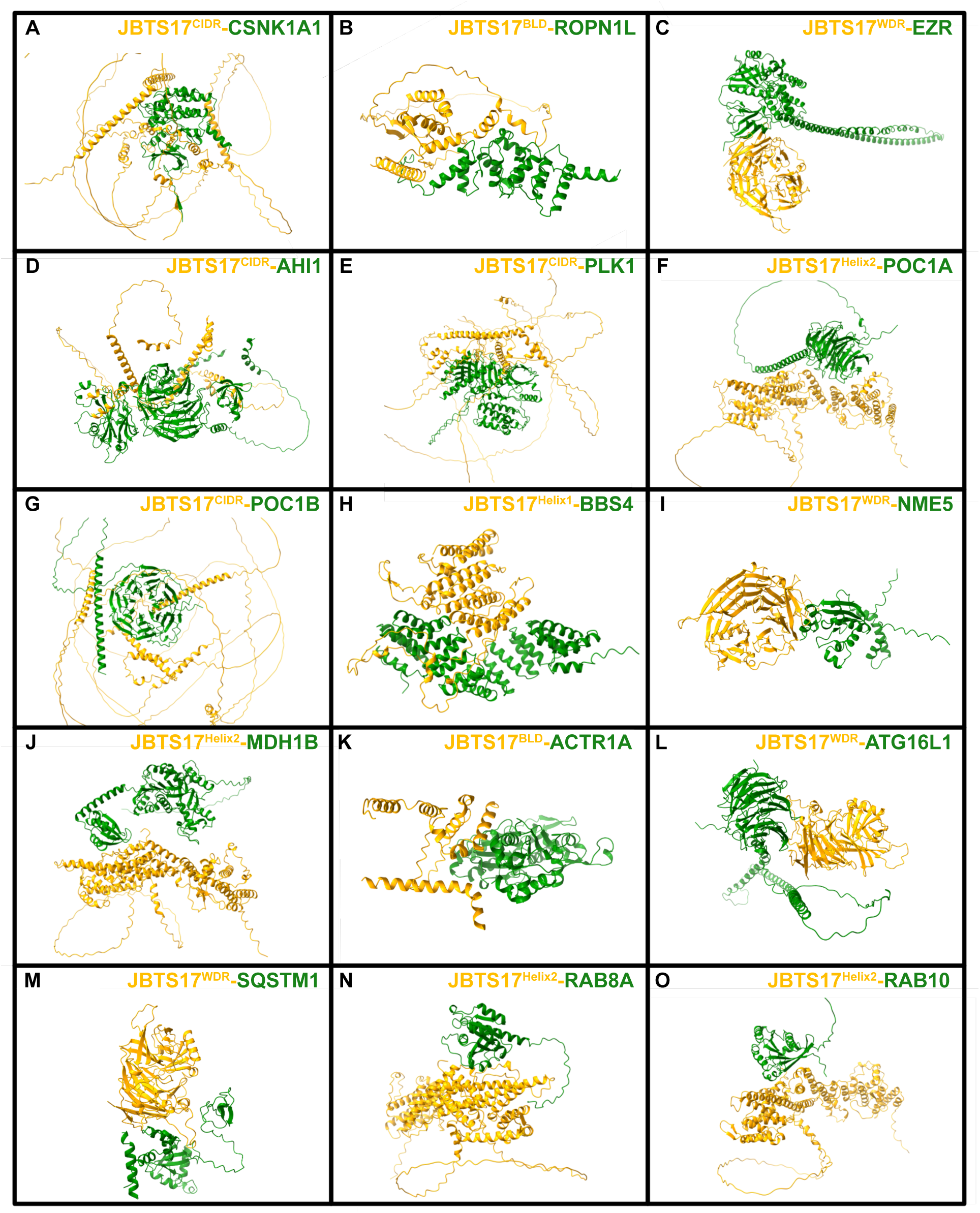
Predicted structures of candidate JBTS17 interactions identified by the AlphaFold screen. AlphaFold2-Multimer-predicted structures of candidate interactions identified in the JBTS17 protein-protein interaction screen and discussed in the text. Each model contains the JBTS17 fragment used in the screen together with the indicated candidate interactor. JBTS17 fragments are shown in gold and candidate proteins are shown in green. (A) JBTS17^CIDR^-CSNK1A1 (B) JBTS17^BLD^-ROPN1L (C) JBTS17^WDR^-EZR (D) JBTS17^CIDR^-AHI1 (E) JBTS17^CIDR^-PLK1 (F) JBTS17^CIDR^-POC1A (G) JBTS17^CIDR^-POC1B (H) JBTS17^Helix1^-BBS4 (I) JBTS17^WDR^-NME5 (J) JBTS17^Helix2^-MDH1B (K) JBTS17^BLD^-ACTR1A (L) JBTS17^WDR^-ATG16L1 (M) JBTS17^WDR^-SQSTM1 (N) JBTS17^Helix2^-RAB8A (O) JBTS17^Helix2^-RAB10

**Supplemental Table 1. Predicted JBTS17 protein interactions and target protein annotations.**

Results of the structure-based interaction screen between five JBTS17 fragments and candidate ciliary proteins. The “All Results” sheet contains the combined screen results, while the “WDR,” “Helix1,” “Helix2,” “BLD,” and “C-IDR” sheets contain results for each individual JBTS17 bait fragment. Each row represents a predicted bait–target interaction and includes the target protein name, primary gene name, interaction confidence scores (ipSAE, ipTM, ipSAE_d0chn, ipSAE_d0dom, ipTM_d0chn, pDockQ, and pDockQ2), and target annotations. The “RFX2 Target,” “Toriyama APMS Hit,” and “CiliaCarta Member” columns indicate whether the target belongs to each respective candidate protein set. The “Annotations” sheet provides additional target information, including protein identifiers, ortholog annotations, protein lengths, predicted disordered regions, and Gene Ontology biological process annotations.

